# Beyond Navigation - Tissue-contacting Fluorescent Lifetime Imaging reveals a pathology-linked lung cancer phenotype at the point of biopsy

**DOI:** 10.64898/2026.08.18.745502

**Authors:** Joel T. Collins, Qiang Wang, Gareth O. S. Williams, Hazel Stewart, Harry A. C. Wood, Charlotte Parry, Caitlin M. Toogood, Annya M. Bruce, Vikki Young, Anne M. Moore, David A. Dorward, Adam D. L. Marshall, Antonella Pellicoro, Lewis Bain, Ahsan R. Akram, Kevin Dhaliwal, James M. Stone

## Abstract

**Background:** Accurate sampling of suspected peripheral lung cancers depends on access to the lesion and confirmation that the biopsy tool is in contact with target tissue. Current bronchoscopic navigation and imaging techniques can guide instruments to a target but do not provide real-time biological confirmation at the point of sampling. Fluorescence lifetime imaging microscopy (FLIM) provides molecular contrast by measuring fluorescence decay - how long photons continue to be emitted from fluorescent molecules. In the Precision Lung clinical study (ISRCTN15093468), the Prothea Imaging System (Generation 1)^i^ identified a candidate tumour-associated phenotype of spatially overlapped low fluorescence lifetime and low intensity (LLLI) from in-vivo imaging. We used this observation as the basis for a reverse-translational study to determine whether the LLLI phenotype is linked to cancer pathology; reproducible with the Imaging System (Generation 2)^ii^; and distinguishable from normal lung tissue.

**Methods:** Previously reported Precision Lung findings were used as the clinical starting observation and were not re-analysed. Validation was then performed using: (i) pathology linked benchtop FLIM of early-stage non-small-cell lung tissue microarrays encompassing malignant cell clusters of approximately 300 µm^2^, matched to the EoT imaging scale; (ii) five sequential fresh lungcancer resections imaged at tumour and comparator regions, including visibly blood-rich contact sites, using the (Generation 2) Imaging System; and (iii) systematic mapping of two ventilated non-cancer donor lungs, one from a smoker and one from a non-smoker, across all available lobes. The LLLI phenotype was defined as spatial co-localisation of low intensity and short lifetime.

**Results:** Using a real time fibre based FLIM system, capable of deployment through a working channel of a bronchoscope, the LLLI tumour phenotype was optically identified in freshly resected tumour tissue. The same phenotype was identified in fixed tissue samples with known pathology, and with images taken in the Precision Lung clinical study. Whole human lung controls did not show evidence of the tumour phenotype.

**Conclusions:** This evidence forms a reverse-translational chain that supports the concept of the Prothea Imaging System - as a platform that confirms that the tool is in contact with a region of cancer in the lesion, while preserving continuous access for biopsy or intervention.

## Introduction

Lung-cancer screening and advances in computed-tomography imaging are increasing the detection of small peripheral pulmonary lesions that require tissue diagnosis [1]. Robotic bronchoscopy, electromagnetic navigation, radial endobronchial ultrasound, and cone-beam computed tomography improve access and geometric localisation, but they do not provide biological confirmation of what is touching the catheter’s distal tip. They can show that a catheter is close to, or within, the radiological target, yet cannot establish in real time that the contact surface contains viable, representative tumour rather than normal lung, inflammation, necrosis or non-lesional tissue. No clinically established bronchoscopic technology currently combines real-time, label-free biological point-of-contact confirmation with a preserved route for biopsy or intervention. This is the beyond-navigation, or final-centimetre, problem.

An effective solution to this problem requires that interpretable data is only acquired when the device is in distal contact with tissue and that the imaged field of view is linked to the deployed biopsy or interventional tool. It should also incorporate an imaging technology capable of adding biological contrast beyond brightness and morphology. Fluorescence Lifetime Imaging Microscopy (FLIM) is suited to the imaging part of this task because it measures the temporal decay of fluorescence following excitation, a parameter influenced by the local tissue biochemical and structural environment. The Eyes on Target (EoT) catheter of the Prothea Imaging System allows direct-contact FLIM in the distal lung whilst maintaining a working channel for image-linked sampling or intervention.

The completed Precision Lung study was designed primarily to establish whether the Generation 1 KronoScan™ (KC.02)/ Eyes on Target (EoT.01) platform could be safely deployed during routine bronchoscopy whilst acquiring FLIM images and permitting tissue sampling through the same catheter. Twenty evaluable participants underwent successful paired imaging and EoT-directed sampling. In the accepted 2026 AABIP conference report, target images were acquired in all evaluable participants, no device-related adverse events were reported, and a significant FLIM-signature change was observed in 94% of malignant lesions [2]. Precision Lung identified the biological pattern that motivated this study. In representative malignant targets, reduced autofluorescence intensity and shortened fluorescence lifetime occurred within the same tissue region, with malignancy confirmed by pathology from EoT-directed sampling. We refer to this spatially clustered state as the co-localised low-lifetime, low-intensity (LLLI), or low-low, phenotype. Low intensity alone may result from poor contact, blood, mucus, attenuation, or low photon yield. A candidate LLLI region is therefore interpretable only when the fluorescence-decay fit is reliably acquired.

Since Precision Lung, Prothea Technologies has developed the next-generation KronoScan™/Eyes on Target (KC.03/EoT.02) system to measure this phenotype more robustly in real time. KC.03 provides eight times more temporal bins at approximately 1.6 times the frame rate, giving more than 12-fold greater temporal-data throughput, together with improved decay estimation, rapid automated calibration, expanded raw-data storage and replay, and lifetime-fit-based quality control.

This study uses a reverse-translational design, testing the in-vivo Precision Lung observation under progressively more controlled conditions. Pathology-linked benchtop FLIM and tumour microarrays first tested the phenotype at approximately 300 µm^2^ scales, including Stage I and Stage IA cancers. Five sequential fresh lung-cancer resections were then imaged with KC.03/EoT.02 at tumour, interface, and comparator regions, including visibly blood-rich contact sites. Finally, ventilated non-cancer donor lungs from a smoker and a non-smoker were systematically mapped to determine whether normal-lung heterogeneity, smoking-related change, ex-vivo ventilation or acquisition conditions reproduce the same persistent clustered phenotype. Together, these experiments tested whether a clinically observed signal can be grounded in pathology and translated into a next-generation platform for biological confirmation at the point of biopsy and, ultimately, intervention.

### Eyes on Target (EoT.02) direct-contact interface

The EoT.02 catheter (Figure 3b) is designed to allow imaging and biopsy sampling in the alveolar space. It is 2.7 m long, has a maximum outer diameter of 1.9 mm and incorporates a working channel of 1.2 mm diameter for the deployment of biopsy tools during visualisation. The catheter embeds an imaging optical fibre within its wall while preserving a 1.2 mm working channel enabling sampling directly adjacent to the image [4]. The proximal end of the catheter is attached to the KronoScan (KC.03) imaging system (Figure 3a), which acquires images of the tissue in direct contact with the catheter’s distal tip. EoT.02 is a prototype device under development and is not yet cleared or approved for clinical use.

**Figure 1.**
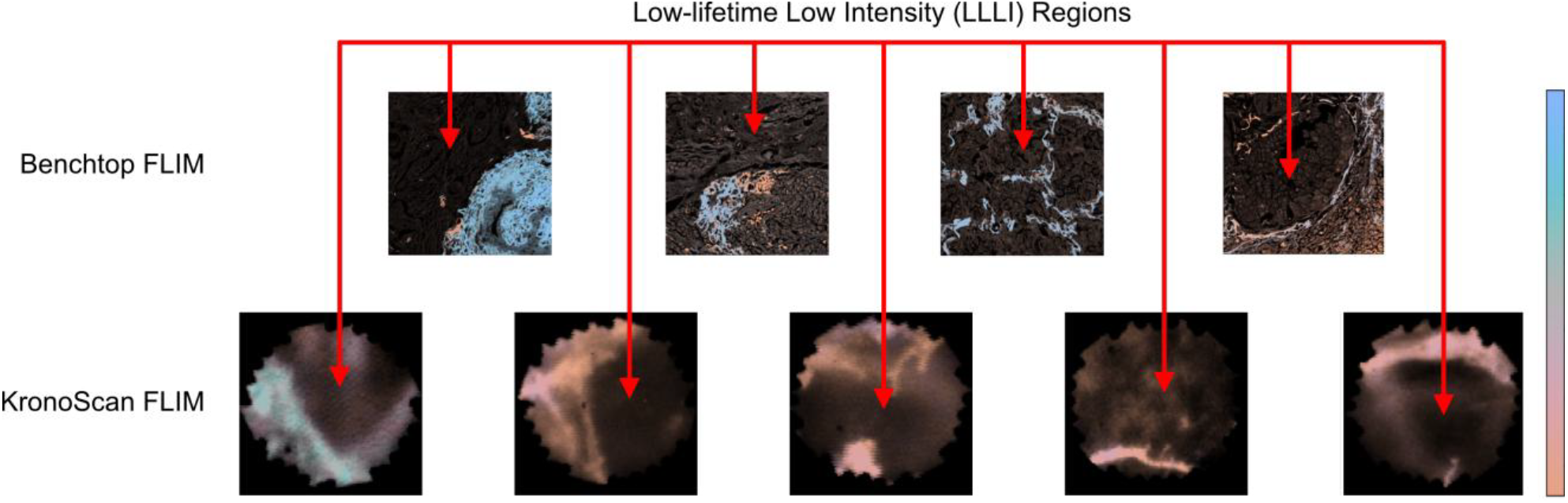
Example frames demonstrating characteristic LLLI phenotype across both benchtop FLIM images and KronoScan FLIM images of cancerous tissue. Cancer subtype, stage, H&E staining confirmation, and colour-map ranges for benchtop FLIM images are provided in Figure 2. Reference tissue comparisons and colour-map ranges for KronoScan FLIM images are provided in Figure 4.

**Figure 2.**
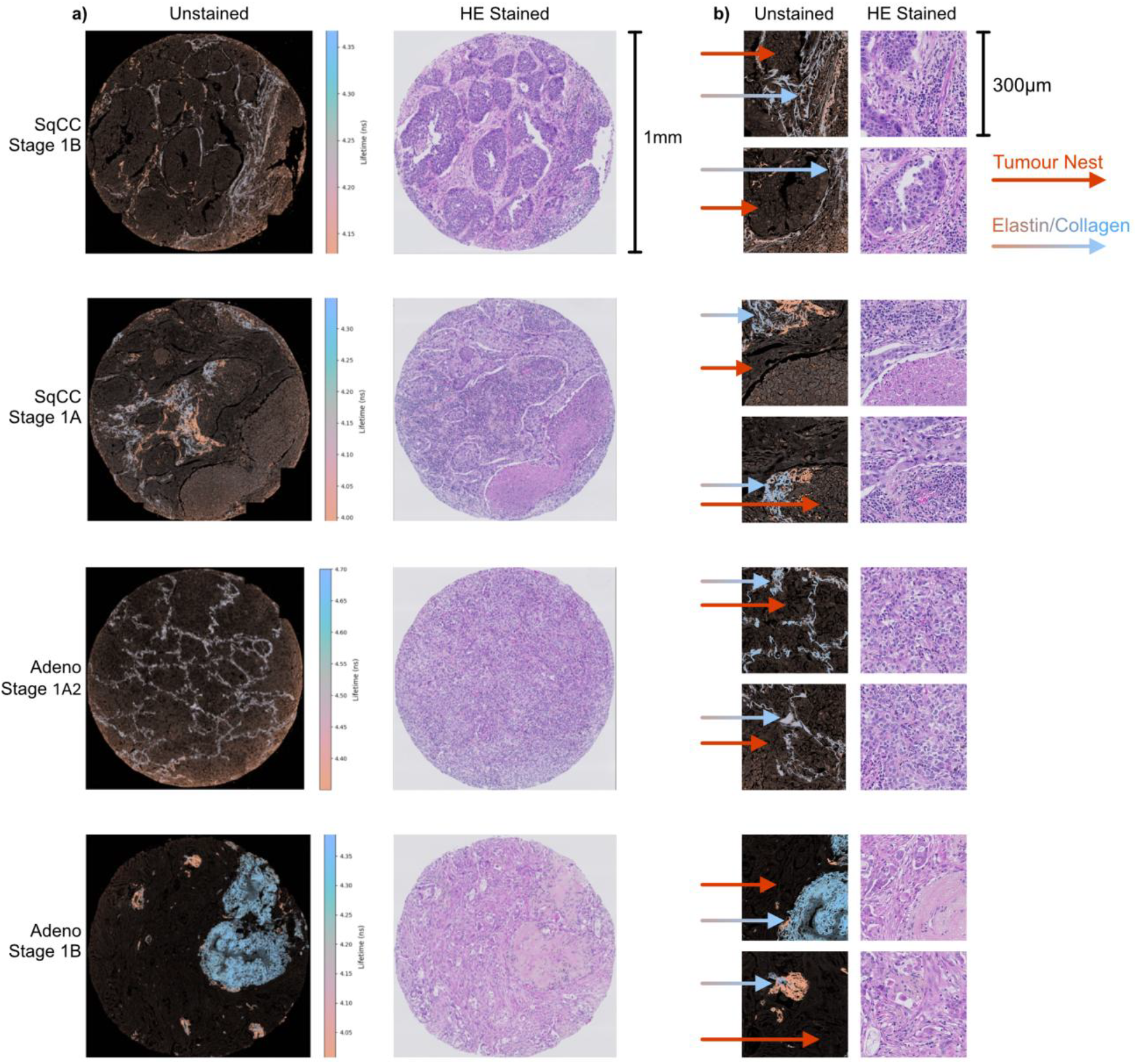
a) Representative lung tumour microarray cores imaged unstained using a benchtop fluorescence-lifetime imaging microscope and subsequently shown with the corresponding H&E-stained sections. Whole-core views demonstrate substantial spatial heterogeneity in the unstained fluorescence appearance across histologically heterogeneous tumour tissue. Scale bar, 1 mm. b) Representative regions from the same cores shown at approximately 300 µm scale, selected to approximate the spatial extent sampled by EoT/KronoScan. Unstained images are paired with the corresponding H&E regions. At this scale, tumour tissue contains broad low-signal areas together with structured brighter features and local fluorescence contrast, closely resembling the spatial appearance observed in the current-generation KronoScan tumour examples shown in Figure 4. Scale bar, 300 µm.

**Figure 3.**
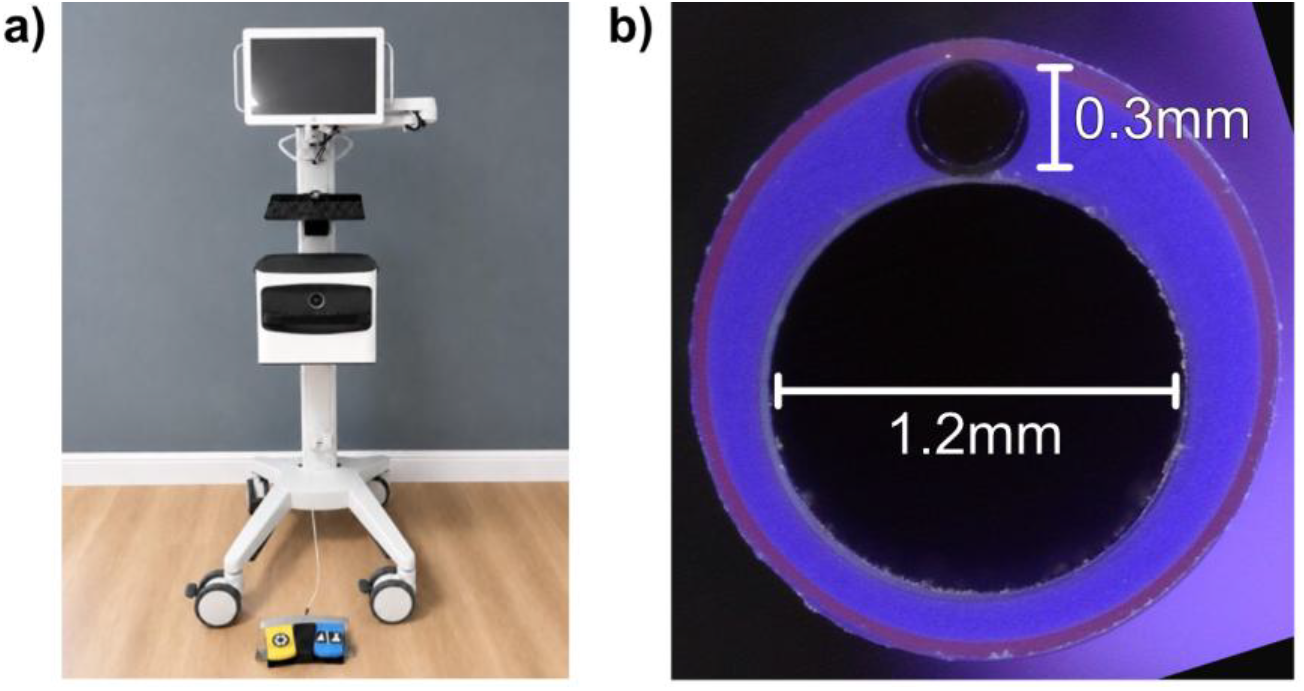
a) KronoScan (KN.03) label-free molecular imaging system b) Distal end of the EoT.02 imaging and access catheter showing the ~0.3mm imaging optical fibre and 1.2mm working channel.

### KronoScan™ Imaging System (KC.03)

KC.03 (Figure 3a) is a confocal fluorescence lifetime imaging platform. The KC.03 prototype system used in this study incorporates eight times more temporal bins and approximately 1.6 times the frame rate of the earlier KC.02 research platform used in the Precision Lung trial, yielding more than 12-fold higher temporal-data throughput. The system supports automated calibration, expanded raw-data storage, real-time replay, and post-acquisition review along with input of annotations to mark different sections of recorded data. These features enable acquisition of intensity and fluorescence-lifetime images with improved temporal sampling and support more advanced lifetime-estimation algorithms than were available on previous versions. The KronoScan KC.03 system represents a substantial engineering evolution from the earlier clinic-ready KronoScan KC.02 research platform. The KC.03 system is a prototype device under development and is not approved for clinical use.

## Results

Figure 1 shows summary frames demonstrating the characteristic LLLI phenotype across both benchtop FLIM images and KronoScan FLIM images of cancerous tissue. The following sections work through the described four-level evidence chain for this reproducible optical cancer phenotype in detail.

### Benchtop FLIM of samples with known pathology

Benchtop fluorescence imaging of lung tumour microarrays (TMAs) demonstrated optical heterogeneity across tumour tissue and provided a pathology-linked reference for interpreting the fibre-based images (Figure 2). At the scale of the complete tissue cores, unstained fluorescence images showed extensive dark regions interrupted by spatially organised brighter structures and localised areas of contrasting fluorescence. Comparison with the corresponding H&E-stained sections demonstrated that this optical heterogeneity occurred within histologically heterogeneous tumour tissue rather than arising from a spatially uniform specimen.

To examine the tissue at a scale more directly comparable with EoT imaging, smaller regions approximately 300 µm across were selected from the unstained TMA images and displayed alongside their corresponding H&E regions. At this scale, the tumour tissue showed a recurring appearance characterised by broad low-signal regions interspersed with structured brighter features, boundaries and localised fluorescence contrast. This appearance closely resembles that observed in the tumour frames acquired with KronoScan in the fresh P2R resections (Figure 4), in which relatively dark tumour regions occur alongside spatially organised fluorescence features.

**Figure 4.**
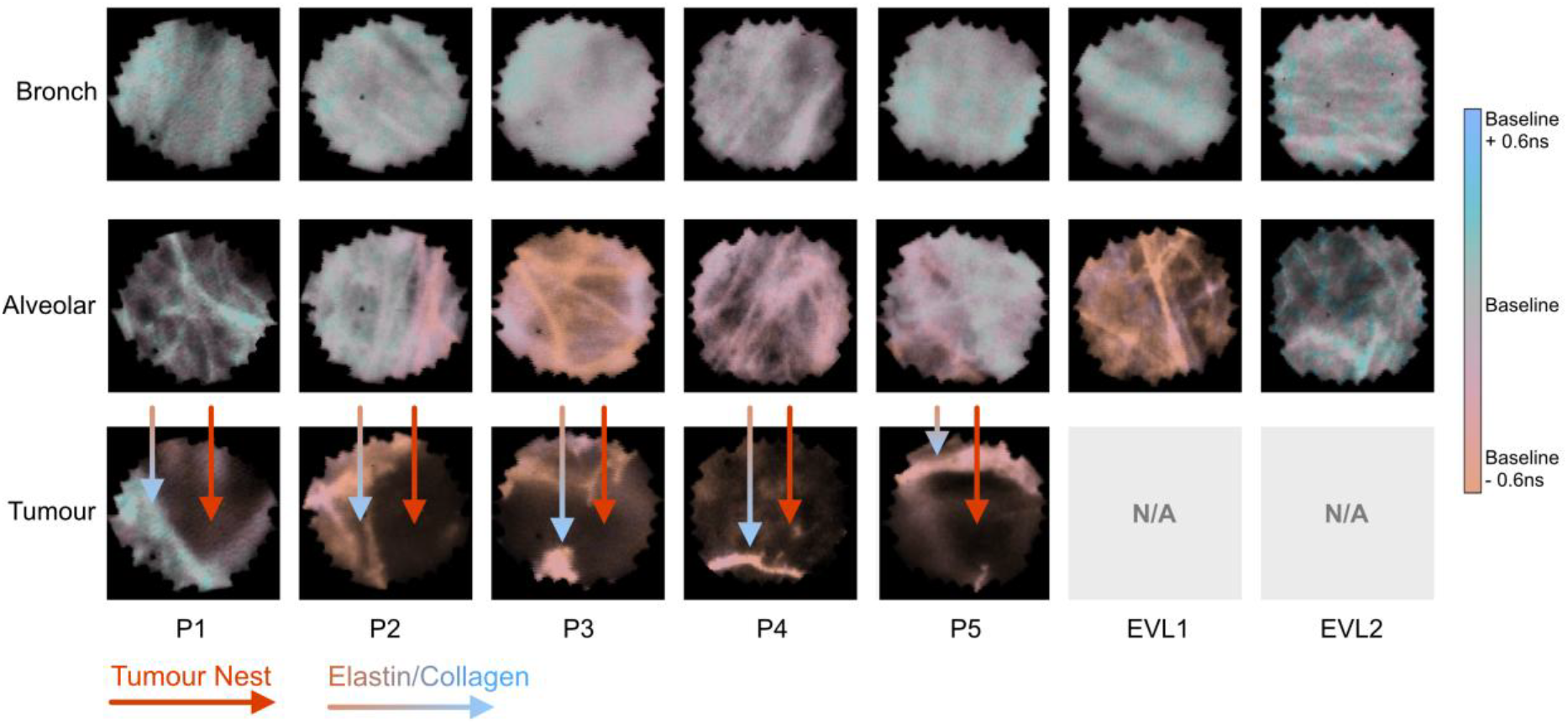
Columns correspond to the five P2R cases and two EVLV procedures. Rows show representative healthy bronchi, healthy alveolar and tumour frames; tumour panels are not applicable to EVL and are shown as blank. Each cell contains the combined (auto-scaled) intensity and (fixed-scale) fluorescence-lifetime representation of one selected frame. The selected P2R tumour frames illustrate spatially organised regions in which reduced intensity co-localises with shortened lifetime (relative to baseline acquired from healthy bronchi tissue), while the within-case bronchi and alveolar frames provide the healthy-tissue visual reference. EVLV panels demonstrate optical heterogeneity in ventilated non-cancer bronchi and alveolar tissue. Representative-frame selection was independent of the case-level quantitative endpoints shown in Figure 6.

The correspondence across spatial scales provides an important bridge between traditional microscopy and optical fibre imaging. The TMA data show that the heterogeneous optical appearance observed with KronoScan can arise from tumour tissue architecture at approximately the same spatial scale sampled by EoT, while the matched H&E sections provide the histological context for those regions. Figure 4 therefore tests whether this pathology-linked appearance is reproduced across independently acquired fresh tumour specimens and distinguishes it from healthy bronchial and alveolar tissue.

### Fresh human lung cancers and whole lung controls using KronoScan™ KC.03 with EoT.02

Figure 3 shows the KronoScan™ KC.03 (a) and EoT.02 (b, c) used in the study.

Representative KC.03 image frames from five P2R fresh-resection cases and two EVL procedures are shown as a case-by-tissue matrix in Figure 4. The rows represent the seven effective cases, while the columns show reference bronchi, reference alveolar and, where available, tumour tissue. The Ex Vivo Lung Ventilation (EVLV) procedures contained no tumour and therefore contribute reference bronchi and alveolar examples only.

Across the five P2R cases, the selected tumour frames contained spatially organised dark regions with corresponding shortening of fluorescence lifetime. Comparison with reference bronchi and alveolar frames from the same case demonstrated that the observation was not based on image darkness alone. Some healthy regions contained low-intensity pixels or local lifetime variation, but the candidate phenotype was identified visually where reduced intensity, and shortened lifetime occupied the same tissue region.

The EVL images demonstrated the range of optical appearances encountered in ventilated non-cancer lung. These images provide reference-tissue context for the P2R examples rather than acting as a binary negative class: healthy bronchi and alveolar tissue could contain dim or relatively shortlived pixels, but the intensity and lifetime channels had to be interpreted together.

Figure 4 is intended to establish the spatial appearance and cross-case reproducibility of the phenotype. The prevalence of the corresponding joint low state was assessed independently from the complete analysed annotation ranges and was not calculated from the selected frames shown in this figure.

To determine whether the representative images reflected the wider P2R and EVLV datasets, a complementary case-level analysis quantified the proportion of usable pixels occupying a joint low-intensity, low-lifetime state.

For each effective P2R or EVL case, the available healthy bronchi and healthy alveolar annotated image ranges were combined to define a within-case reference distribution. Intensity was expressed relative to the median reference intensity. The 25th percentiles of the reference intensity and lifetime distributions were then used as case-specific lower-tail cut-offs. The joint low-state fraction was defined as the percentage of usable pixels falling below both cut-offs.

Each annotated image range was summarised independently, and tissue-level values were calculated by averaging the image-range fractions. This prevented a long recording or a range containing many repeated frames from receiving greater weight solely because it contributed more pixels.

In all five P2R cases, tumour tissue had a larger joint low-state fraction than healthy bronchi from the same case (Figure 6). The available tumour-versus-healthy-alveolar comparisons were also positive. The magnitude of the effect varied between cases, consistent with biological and acquisition heterogeneity, but the within-case direction was in agreement with the image-level pattern shown in Figure 4.

**Figure 5.**
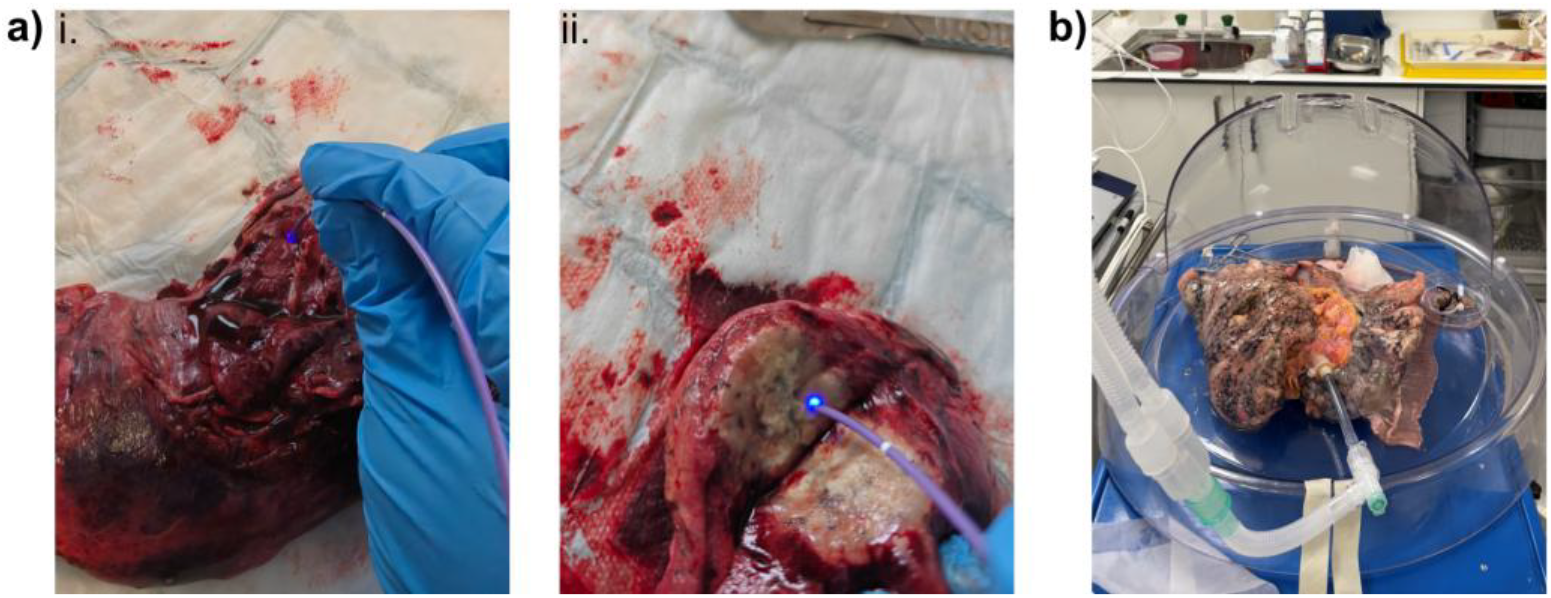
a) Direct contact with grossly identified tumour-rich regions of fresh, visibly blood-rich lung cancer resections, showing contact with reference tissue (i), and tumour tissue (ii). b) Image of ventilated smoker’s lung used for mapping in EVL1.

**Figure 6.**
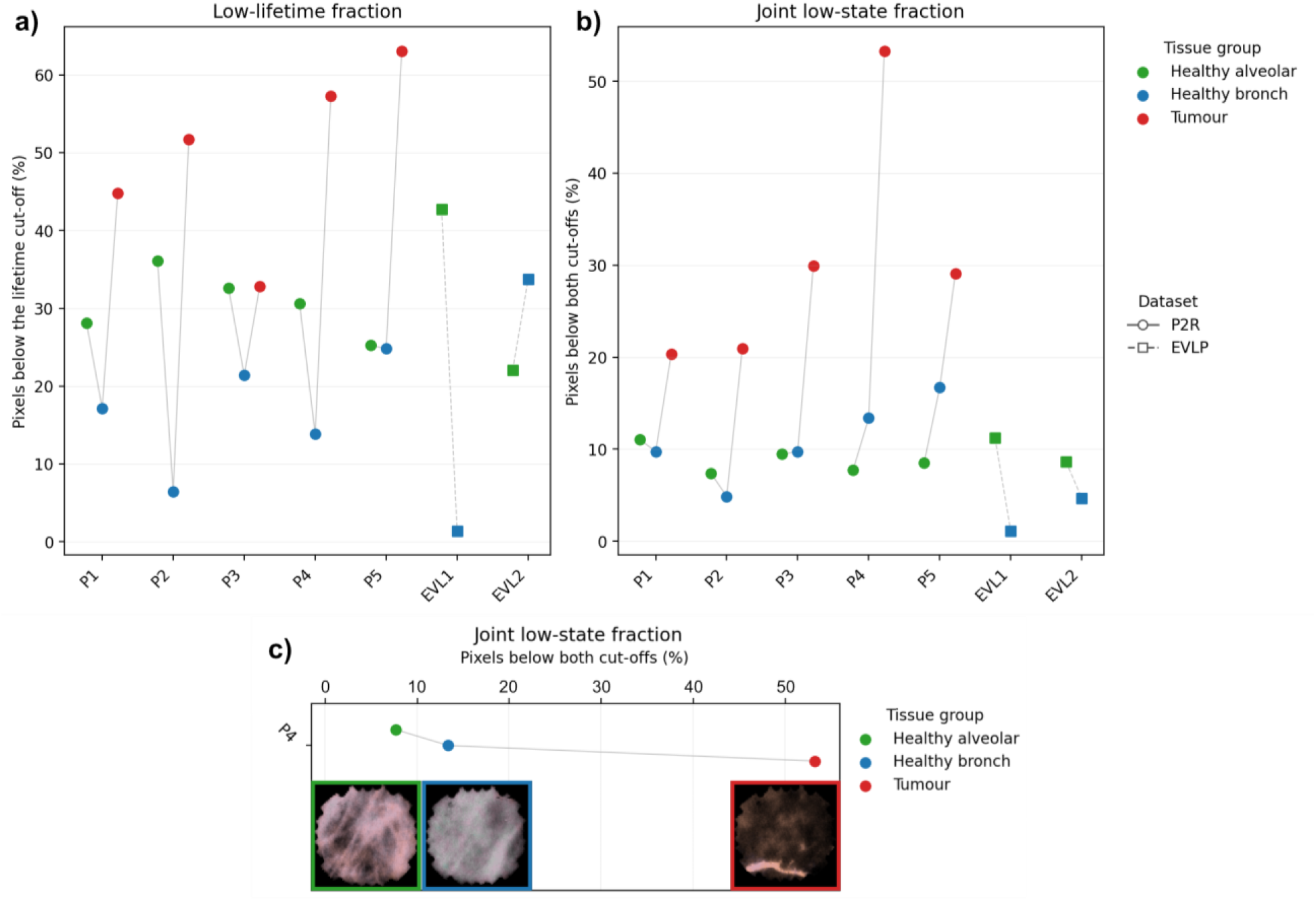
a) Low-lifetime fraction, defined as the percentage of usable pixels below the case-specific lifetime cut-off (25th percentile of the healthy-reference distribution) b) Joint low-state fraction relative to healthy bronchi from the same cases. The joint low state was defined as the percentage of usable pixels below both the case-specific intensity and lifetime cut-offs (25th percentile of the healthy-reference distribution). Positive values indicate a larger joint low-state fraction. Healthy-reference distributions were obtained by pooling both reference bronchi and alveolar data. c) Joint low-state fraction alongside example frames from Figure 4 for case P4.

The bronchi-to-alveolar relationship was more variable across the P2R and EVLV cases. EVLV contributed healthy-tissue comparisons only and was not assigned a tumour effect. These healthytissue data demonstrate that lower-tail pixels are not exclusive to tumour and that the relevant quantitative result is an increased joint low-state fraction relative to the corresponding within-case reference.

The joint-state endpoint is a distributional summary. It supports recurrence of the optical pattern across effective cases but does not retain information about cluster shape, spatial location or persistence between consecutive frames.

### Precision Lung agreement

Precision Lung is included as the clinical starting point for this reverse-translational study. In the completed prospective study, 20 evaluable patients were successfully bronchoscoped with paired target and non-lesion imaging, and sampling was performed through the integrated EoT channel. The accepted AABIP report described target image acquisition in all evaluable cases, no device-related adverse events and a significant FLIM-signature change in 94% of malignant lesions[2].

Figure 7 shows the representative pathology-linked example previously reported from a participant. Compared with the paired control field, the lesion contained lower-intensity regions and a shift to shorter lifetime. Tissue obtained through EoT from the imaged target confirmed squamous-cell carcinoma on H&E histology. This example illustrates the clinical observation that prompted subsequent pathology-scale, fresh-resection and ventilated-lung experiments.

**Figure 7.**
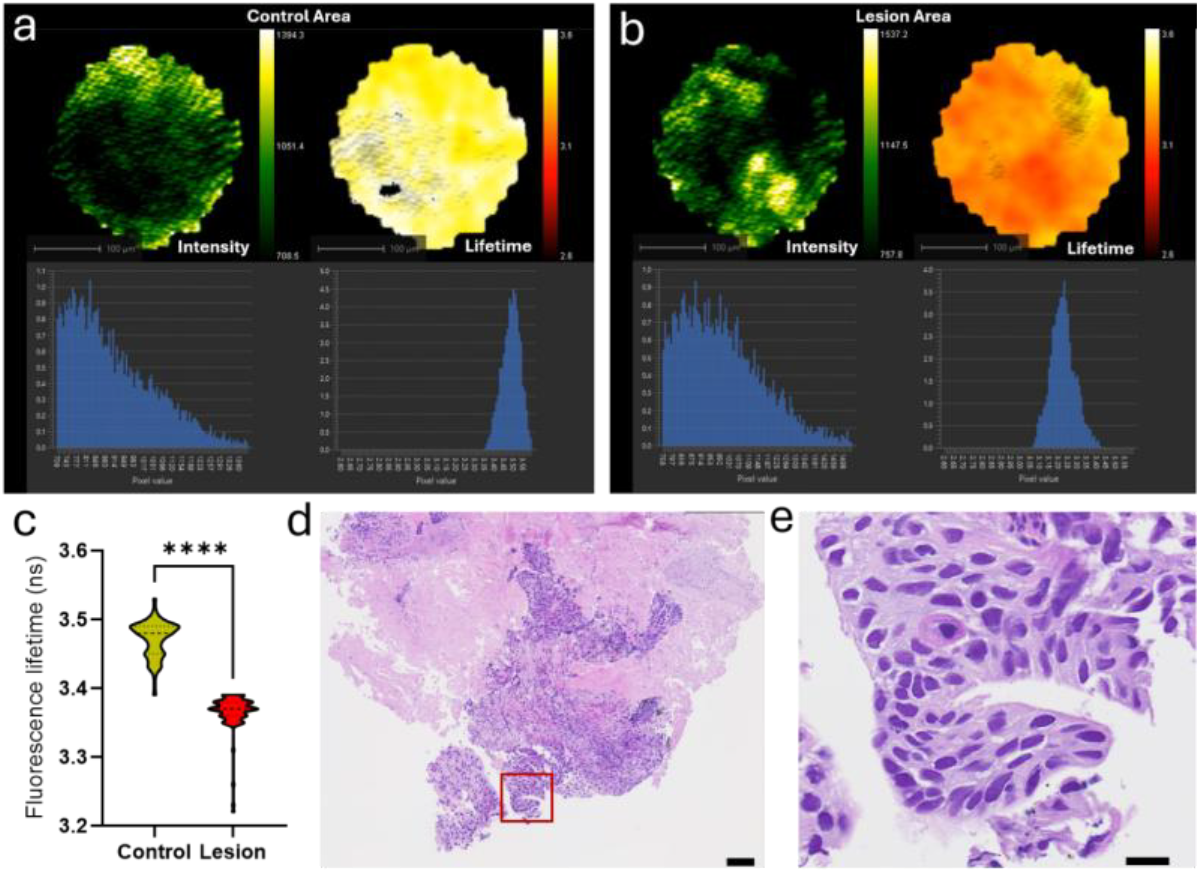
Previously reported in-vivo Precision Lung example that motivated the reverse-translational validation programme. Representative intensity and lifetime images are shown for a paired control region and lesion region, with pixel distributions. The lesion contains lower-intensity regions and shorter lifetime than the paired control. EoT-directed biopsy from the imaged target confirmed squamous-cell carcinoma on H&E histology. The figure is included as previously communicated visual concordance from the accepted AABIP report; the Precision Lung cohort is not re-analysed for diagnostic performance in the present manuscript. Scale bars as shown in the source figure.

Precision Lung used the earlier KC.02 acquisition and lifetime-estimation workflow. The present paper therefore treats its representative images as directional in-vivo concordance and does not pool numerical thresholds or effect sizes across system generations. Clinical decisions were not made from FLIM in Precision Lung, and prospective biopsy-guidance validation remains required.

## Discussion

It has been shown that low-intensity regions are common in optical endomicroscopy of tissue and may arise from biological structure, variable photon collection, obscuration, or imaging conditions. Intensity alone therefore cannot determine why a region appears dark. Traditional benchtop FLIM, free from the constraints of fibre-based imaging, confirms that there are genuine low intensity regions within tumour tissue samples and that these generally correspond to dense cancerous regions with reduced lifetime.

In this study, tumour regions imaging via the KC and EoT endomiscopscopy platform were characterised by the co-localisation of reduced fluorescence intensity and shortened fluorescence lifetime as an optical phenotype for cancer. Ventilated lungs with no evidence of cancer were used as a control and demonstrated that low intensity can occur in the absence of the corresponding lifetime change, and vice versa, but with limited overlap of the two signal types. This supports the interpretation that the optical cancer phenotype requires a spatial convergence of reduced intensity and shortened lifetime, which discriminates against artifacts related to loss of contact with the sample.

The KC and EoT platform acquires images through direct tissue contact rather than a defined optical working distance. This allows for interpretable intensity and lifetime measurements to be obtained from fresh surgical specimens, despite visible blood at multiple imaging sites. The integrated working channel enables tissue sampling adjacent to the imaging field, preserving the spatial relationship between optical assessment and biopsy. Together, these features support the practical use of fluorescence lifetime imaging at the point of tissue sampling.

This study was designed to establish the reproducibility of a qualitative optical phenotype rather than a diagnostic classifier. The cancer cohort was small, and additional studies are required to assess performance across larger populations and clinically relevant benign comparators, including inflammation, fibrosis, necrosis, and treatment-related change. Future work should also examine blinded interpretation, interobserver agreement, and pathology-linked quantitative definitions of the phenotype.

## Conclusions

This study closes a reverse-translational loop that began in patients. In the completed Precision Lung study, 20 evaluable participants underwent successful bronchoscopic EoT imaging, and the accepted AABIP report described malignant targets containing lower-intensity and shorter-lifetime regions. In this study, that in-vivo observation was correlated to pathology, reproduced with the current-generation KronoScan™/EoT platform in all five fresh human lung cancers examined, and tested against ventilated non-cancer whole-lung controls from a smoker and a non-smoker.

The candidate optical phenotype is not image darkness alone. It is the spatial overlap of reduced intensity and shortened lifetime. Its presence in pathology-confirmed early-stage cancers, including Stage IA disease, supports relevance to the small lesions increasingly detected through screening. Its non-reproduction as a persistent clustered state in ventilated non-cancer lungs, together with successful acquisition at visibly blood-rich resection sites in resected tissue, strengthens the interpretation that it reflects tumour-associated biology rather than a generic imaging artefact.

Direct-contact FLIM therefore provides a route to pathology-linked optical confirmation at the point of biopsy while maintaining a 1.2 mm path for tissue acquisition and intervention. The platform remains investigational.

## Methods

### Pathology-resolved reference images

Early-stage non-small cell lung carcinoma (NSCLC) tissues were obtained from surgically resected specimens archived in a clinical biobank and processed using a standard formalin-fixed, paraffin-embedded (FFPE) workflow, approved by the NHS Lothian NRS Bioresource (REC No. 20/ES/0061), with a specific application SR1949. Tissue sections of 4 μm thickness were prepared and mounted onto glass slides. Prior to imaging, the sections were deparaffinized in xylene and subsequently rehydrated through a graded series of ethanol solutions. A cohort of four early-stage NSCLC patients (Stage I) were used as the reference images, including two adenocarcinoma and two squamous cell carcinoma cases. Afterwards, unstained samples were imaged using a confocal FLIM system (Leica STELLARIS 8 FALCON FLIM Microscope) with a 20x/0.75NA objective, using excitation and emission wavelengths at 488 nm and [500 nm, 720 nm]. Lifetime images were reconstructed from the raw data using the 4-exponential fitting algorithm in Leica LAX-X software. The unstained tissue was then Haematoxylin and Eosin (H&E) stained as the gold standard of the corresponding FLIM images.

FLIM datasets were processed using the workflow described in previous studies [5], except for the visualization procedure. To maintain consistency and facilitate direct visual comparison with fibre-based FLIM images, the mean tumour lifetime measured from the reference FLIM images was first determined and used as the baseline value for visualisation. For each image patch extracted from the original reference FLIM dataset, the upper bound of the colour scale was adaptively defined according to the maximum lifetime observed within that specific patch.

### Fresh-resection specimens and imaging workflow

Five fresh lung-cancer resections were examined after surgical removal. Material provision and use was covered by Lothian NRS BioResource RTB approval (REC ref – 25/ES/0030). Tumour, non-tumour and, where available, tumour margin, tissue regions identified before imaging. The tumour tissue was sliced open to ensure good contact of the distal tip of the EoT.02 catheter and the target tissue (Figure 5a). The tissue samples remained in their fresh surgical state with visible blood surrounding multiple contact sites. The presence of blood did not inhibit imaging.

All five fresh specimens were surgically resected lung cancers confirmed by routine histopathology.

### Ex-vivo Ventilated whole-lung negative controls (EVLV)

Human donor lungs declined for transplantation were procured from either the International Institute for the Advancement of Medicine in the United States following approval from the appropriate regional ethics committee and with informed consent from donors or their next of kin or from the UK (20/ES/0061). Two non-cancer whole human lungs (one smoker, both left and right lung (“20260605 EVLV”), and one non-smoker, right lung only (“20260723 EVLV”)) were maintained under ex vivo ventilation using a Drager Savina 300 ventilator with a tidal volume of 3-6 mL/Kg ideal body weight, positive end-expiratory pressure of 5cm H^2^O, pressure support of 5cm H^2^O, and respiratory rate of 12 breaths/min and systematically mapped with KC.03 system. Imaging was recorded across all lobes and subsegments of all lungs as a negative control.

### Precision Lung Images

This manuscript treats representative images from the Precision Lung Study (ISRCTN15093468 [2]) as previously collected *in-vivo* evidence of visual agreement of the LLLI phenotype across ex vivo and does not re-analyse the clinical cohort, safety outcomes or diagnostic-yield data (Figure 7).

### Image-quality gating and processing

For each image pixel, the temporal histogram at each bin was spatially box-filtered using a 7 × 7 kernel before lifetime estimation. Intensity was calculated separately from the unsmoothed histograms. Additional Gaussian filtering was applied when producing the live or replayed display images, with separate configured widths for intensity and lifetime. These filtering operations were preprocessing or visualisation steps and were not themselves lifetime-quality acceptance criteria.

Lifetime validity was evaluated on a per-pixel basis. A lifetime value is retained when the integrated histogram intensity is at least 100 counts, the estimated lifetime is finite and positive, and its relative standard error is no greater than 0.1. Pixels that fail these criteria are assigned no valid lifetime value and are excluded from lifetime-based analyses. These criteria assess photon statistics and numerical estimation quality; they do not include automatic classification of tissue contact, motion, blood, mucus or other field contents.

### Operational definition of the LLLI optical phenotype

The LLLI optical phenotype is an image-level observation in which a region of reduced fluorescence intensity is accompanied by a corresponding reduction in fluorescence lifetime within the same tissue area. The key feature is the spatial co-localisation of these two optical characteristics rather than a decrease in either measurement alone.

In this study, the phenotype was assessed qualitatively from matched intensity and lifetime images acquired under direct tissue contact. Regions of interest were interpreted in the context of adjacent reference tissue from the same specimen or imaging session. Low intensity alone was not considered sufficient evidence of the phenotype, as reduced intensity can arise from multiple biological or technical factors. Fluorescence lifetime provided the additional contrast required to distinguish biologically altered dim tissue from regions that were simply less fluorescent.

### Statistical approach

The primary quantitative endpoint was the joint low-state fraction, defined as the percentage of usable pixels falling below both the case-specific fluorescence-intensity and fluorescence-lifetime cut-offs. The image-defined phenotype was assessed separately from representative intensity and lifetime frames. The joint low-state fraction was used as a supportive distributional summary and did not measure spatial connectedness, cluster morphology or persistence across frames.

For the P2R and EVLV datasets, comparisons were made at the case level. Within each case, the available reference bronchi and alveolar annotated image ranges were combined to define the reference distribution. Case-specific 25th-percentile intensity and lifetime cut-offs were then applied to all tissue groups from that case. Each labelled annotated image range was summarised separately, and tissue-level estimates were calculated by averaging the image-range fractions. This prevented longer recordings or ranges containing more frames from receiving greater weight solely because they contributed more pixels.

Low-lifetime and joint low-state fractions were displayed for healthy bronchi, healthy alveolar and tumour tissue, where annotated. Comparisons were made descriptively between tissue groups from the same case. EVLV procedures contained healthy bronchi and alveolar annotated images only and therefore provided healthy-tissue context rather than tumour comparisons.

Precision Lung was analysed separately because it was acquired and processed using the earlier KronoScan KC.02 workflow. For the 20 complete cases, the saved control distribution defined the case-specific intensity and lifetime cut-offs, which were then applied to the corresponding imaging data. Joint low-state fractions were reported for the paired control and experimental groups.

Pixels and frames were treated as nested measurements within annotated image ranges, contacts, and cases rather than as independent biological replicates. Spatial preprocessing also introduced correlation between neighbouring pixels. Pixel counts were therefore used to estimate within-case distributions but did not determine the biological sample size. The KC.03 and KC.02 Precision Lung results were not pooled because their acquisition systems, lifetime-estimation methods, reference groups and aggregation procedures differed. Given the small number of effective P2R and EVL cases, the analyses were descriptive and emphasised the consistency, direction, and between-case variation of the paired tissue-group fractions.

## Declarations

### Ethics and tissue governance

Images from the Precision Lung Clinical Study were sourced from a poster presented at the 2026 AABIP Conference (Dr Adam Marshall, Principal Investigator) and were collected as part of an approved clinical study conducted in Edinburgh, UK (ISRCTN15093468). No additional participant-level clinical analysis was performed for this manuscript.

For ex vivo tissue analysis, each patient provided written informed consent, and the proposal was approved by Lothian NRS BioResource RTB approval (Regional Ethics Committee reference – 25/ES/0030).

Whole explanted lungs were obtained under ethical approval 20/ES/0061 (UK) and International Advancement of Medicine (US),.

Approval for the fixed tissue samples was obtained under the NHS Lothian NRS Bioresource (REC No. 20/ES/0061).

### Data availability

De-identified source intensity, photon-arrival, lifetime, quality-control and pathology-annotation data should be made available according to institutional governance and commercial/IP constraints. The final statement should specify the repository or controlled-access process.

## Acknowledgements

The authors thank the thoracic surgical, pathology, bronchoscopy, research and engineering teams who supported tissue acquisition and imaging and the Precision Lung Clinical Research Team

## Footnotes

i KronoScan™ (KC.02)/Eyes on Target (EoT.01)

ii KronoScan™ (KC.03)/Eyes on Target (EoT.02)

## References

[1] Y. Ru Zhao, X. Xie, H. J. de Koning, W. P. Mali, R. Vliegenthart and M. Oudkerk, “NELSON lung cancer screening study,” Cancer Imaging, vol. 11, p. S79–S84, 2011.

[2] A. D. L. Marshall, F. R. Millar, D. R. Dorward, A. R. Akram, S. Giavedoni, L. Bain, K. Hamilton, J. Antonelli, B. Wheeler, A. Bruce, H. Stewart, J. Collins, G. Williams, J. Stone, N. A. Hirani and K. Dhaliwal, “Fibre-Based Fluorescence Lifetime Imaging (FLIM) of Peripheral Lung Lesions: A First in Human Study,” in 9th Annual Conference of the American Association for Bronchology and Interventional Pulmonology (AABIP), 2026.

[3] M. Wang, F. Tang, X. Pan, L. Yao, X. Wang, Y. Jing, J. Ma, G. Wang and L. Mi, “Rapid diagnosis and intraoperative margin assessment of human lung cancer with fluorescence lifetime imaging microscopy,” BBA Clinical, vol. 8, p. 7–13, December 2017.

[4] J. M. Stone, H. A. C. Wood, K. Harrington and T. A. Birks, “Low index contrast imaging fibers,” Optics Letters, vol. 42, p. 1484, April 2017.

[5] Q. Wang, S. Fernandes, G. O. S. Williams, N. Finlayson, A. R. Akram, K. Dhaliwal, J. R. Hopgood and M. Vallejo, “Deep learning-assisted co-registration of full-spectral autofluorescence lifetime microscopic images with H&E-stained histology images,” Communications Biology, vol. 5, October 2022.

[6] A. C. Adams, A. Kufcsák, C. Lochenie, M. Khadem, A. R. Akram, K. Dhaliwal and S. Seth, “Fibre-optic based exploration of lung cancer autofluorescence using spectral fluorescence lifetime,” Biomedical Optics Express, vol. 15, p. 1132, January 2024.

[7] R. Datta, T. M. Heaster, J. T. Sharick, A. A. Gillette and M. C. Skala, “Fluorescence lifetime imaging microscopy: fundamentals and advances in instrumentation, analysis, and applications,” Journal of Biomedical Optics, vol. 25, p. 1, May 2020.

[8] S. Fernandes, E. Williams, N. Finlayson, H. Stewart, C. Dhaliwal, D. A. Dorward, W. A. Wallace, A. R. Akram, J. Stone, K. Dhaliwal and G. O. S. Williams, “Fibre-based fluorescence-lifetime imaging microscopy: a real-time biopsy guidance tool for suspected lung cancer,” Translational Lung Cancer Research, vol. 13, p. 355–361, February 2024.

[9] T. Kramer, L. Wijmans, M. de Bruin, T. van Leeuwen, T. Radonic, P. Bonta and J. T. Annema, “Bronchoscopic needle-based confocal laser endomicroscopy (nCLE) as a real-time detection tool for peripheral lung cancer,” Thorax, vol. 77, p. 370–377, June 2021.

[10] C. J. Manley, T. Kramer, R. Kumar, Y. Gong, H. Ehya, E. Ross, P. I. Bonta and J. T. Annema, “Robotic bronchoscopic needle-based confocal laser endomicroscopy to diagnose peripheral lung nodules,” Respirology, vol. 28, p. 475–483, December 2022.

[11] S. Ranjit, L. Malacrida, D. M. Jameson and E. Gratton, “Fit-free analysis of fluorescence lifetime imaging data using the phasor approach,” Nature Protocols, vol. 13, p. 1979–2004, September 2018.

[12] S. Seth, A. R. Akram, P. McCool, J. Westerfeld, D. Wilson, S. McLaughlin, K. Dhaliwal and C. K. I. Williams, “Assessing the utility of autofluorescence-based pulmonary optical endomicroscopy to predict the malignant potential of solitary pulmonary nodules in humans,” Scientific Reports, vol. 6, August 2016.

[13] L. Thiberville, S. Moreno-Swirc, T. Vercauteren, E. Peltier, C. Cavé and G. Bourg Heckly, “In Vivo Imaging of the Bronchial Wall Microstructure Using Fibered Confocal Fluorescence Microscopy,” American Journal of Respiratory and Critical Care Medicine, vol. 175, p. 22–31, January 2007.

[14] Q. Wang, J. R. Hopgood, S. Fernandes, N. Finlayson, G. O. S. Williams, A. R. Akram, K. Dhaliwal and M. Vallejo, “A layer-level multi-scale architecture for lung cancer classification with fluorescence lifetime imaging endomicroscopy,” Neural Computing and Applications, vol. 34, p. 18881–18894, June 2022.

[15] G. O. S. Williams, E. Williams, N. Finlayson, A. T. Erdogan, Q. Wang, S. Fernandes, A. R. Akram, K. Dhaliwal, R. K. Henderson, J. M. Girkin and M. Bradley, “Full spectrum fluorescence lifetime imaging with 0.5 nm spectral and 50 ps temporal resolution,” Nature Communications, vol. 12, November 2021.

[16] Q. Wang, A. R. Akram, D. A. Dorward, S. Talas, B. Monks, C. Thum, J. R. Hopgood, M. Javidi and M. Vallejo, “Deep learning-based virtual H – E staining from label-free autofluorescence lifetime images,” npj Imaging, vol. 2, June 2024.

